# LFAQ: towards unbiased label-free absolute protein quantification by predicting peptide quantitative factors

**DOI:** 10.1101/328864

**Authors:** Cheng Chang, Zhiqiang Gao, Wantao Ying, Yan Zhao, Yan Fu, Songfeng Wu, Mengjie Li, Guibin Wang, Xiaohong Qian, Yunping Zhu, Fuchu He

**Author notes:** Correspondence should be addressed to the following: Yan Fu, Xiaohong Qian, Yunping Zhu, Fuchu He. These authors contributed equally to this work (Cheng Chang, Zhiqiang Gao, Wantao Ying).

## Abstract

Mass spectrometry (MS) has become a prominent choice for large-scale absolute protein quantification, but its quantification accuracy still has substantial room for improvement. A crucial issue is the bias between the peptide MS intensity and the actual peptide abundance, i.e., the fact that peptides with equal abundance may have different MS intensities. This bias is mainly caused by the diverse physicochemical properties of peptides. Here, we propose a novel algorithm for label-free absolute protein quantification, LFAQ, which can correct the biased MS intensities by using the predicted peptide quantitative factors for all identified peptides. When validated on datasets produced by different MS instruments and data acquisition modes, LFAQ presented accuracy and precision superior to those of existing methods. In particular, it reduced the quantification error by an average of 46% for low-abundance proteins.

## Introduction

The quantitative exploration of biological systems usually requires accurate descriptions of protein abundances and their changes on a proteome-wide scale. Mass spectrometry (MS)-based quantitative proteomics has become an important research area of life science^1^. There are two main types of strategies in quantitative proteomics, i.e., relative and absolute protein quantification. Relative protein quantification aims to compare the changes in abundance of the same protein between different samples, whereas absolute protein quantification focuses on determining the abundance (e.g., copy number or molar concentration) of each protein in the same sample^2^. Although highly demanded, large-scale absolute protein quantification remains a challenging task^3^.

Ideally, absolute quantification requires a labeled internal standard (such as QconCAT^4^ or PSAQ^5^) for every protein of interest^6-8^. For the large-scale absolute quantification of complex samples (e.g., an entire proteome of cells, tissues or body fluids), a practical compromise is to use a few spike-in standard proteins to fit a linear function between the calculated MS intensities and the actual abundances of the standard proteins and then to use this function to extrapolate the abundances of all the identified proteins in the sample based on their calculated MS intensities^9^.

However, the raw MS intensity of a peptide is actually a biased measurement of its actual abundance^10, 11^, and this bias can be propagated from the peptide level to the protein level, leading to inaccurate protein quantification. In essence, apart from the matrix effect, the bias in peptide MS intensity is intrinsically caused by the MS behavior of a peptide, which is dominated by the synergistic effect of various physicochemical properties of the peptide^10^. The MS intensity bias is hard to model and often ignored by absolute quantification algorithms^12, 13^. For example, the commonly used algorithms Top3^12^ and iBAQ^13^ employ the unprocessed MS intensities of identified peptides to calculate the protein MS intensities. Although such algorithms are effective for proteins with many identified peptides, they may become inaccurate for small or low-abundance proteins, in which only a few peptides are usually identified.

To eliminate the quantification bias or select representative peptides for accurate large-scale protein quantification, a parameter called peptide detectability^14^ (defined as the probability of a peptide being identified by MS) was introduced in previous studies^15-17^. For example, Jarnuczak *et al.* defined the peptide “flyability” factor to represent the intrinsic peptide detectability in terms of selecting high-responding peptides in targeted MS studies^15^, and APEX^17^ applied peptide detectability to correct protein spectral counts for absolute protein quantification.

Although these studies have made significant progress towards accurate protein quantification, peptide detectability is essentially unsuitable for reducing the MS intensity bias because it represents the probability that a peptide can be identified rather than the extent to which it can be quantified. In fact, peptide detectability has mainly been used to predict proteotypic peptides^18^ or quantotypic peptides^8^ in targeted proteomic applications, in which only a couple of selected peptides are used to quantify each protein of interest. While, the unselected part (often majority) of identified peptides, which may potentially contain important quantification information, are neglected. Moreover, the peptide detectability used in protein quantification^15, 16^ is usually determined from protein or peptide standards, which limits its application in complex biological samples. In summary, there is yet no suitable algorithm equipped with a standard-free machine learning technique to directly correct the MS intensities of all identified peptides for accurate protein quantification.

Here, we present a label-free absolute protein quantification algorithm (LFAQ) that can correct the biased peptide MS intensities using a novel parameter termed the peptide quantitative factor (Q-factor for short). The Q-factor of a peptide reflects the extent to which peptide abundance can be expressed by its raw MS intensity and can be predicted empirically from the physicochemical properties of the peptide without training on standards or manually curated data. We have demonstrated the advantages of LFAQ over previous methods on both our own and public MS data produced from complex samples in data-dependent acquisition (DDA) and data-independent acquisition (DIA) modes. The results showed that the correction of raw MS intensity for each identified peptide using the Q-factor resulted in more accurate and precise quantifications in all experiments. Finally, we provide an easy-to-use software tool to facilitate the application of the LFAQ algorithm.

## Results

### LFAQ algorithm

LFAQ is proposed to accurately calculate protein MS intensity by reducing the biases of the raw MS intensities of peptides using predicted peptide Q-factors. The LFAQ algorithm is shown in **Fig. 1**. First, the peptide Q-factor prediction model is trained by a machine learning approach. It takes the sequences and MS intensities of the identified peptides as input to generate the training set of peptide Q-factors. Then, the Q-factor of each identified peptide in the sample is predicted using the trained model and used to correct the peptide MS intensity. Finally, the LFAQ value (the MS intensity of each protein calculated using the LFAQ algorithm) is obtained by averaging the corrected intensities of all the identified peptides of the protein as follows:

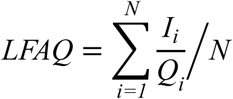

**Figure 1.**
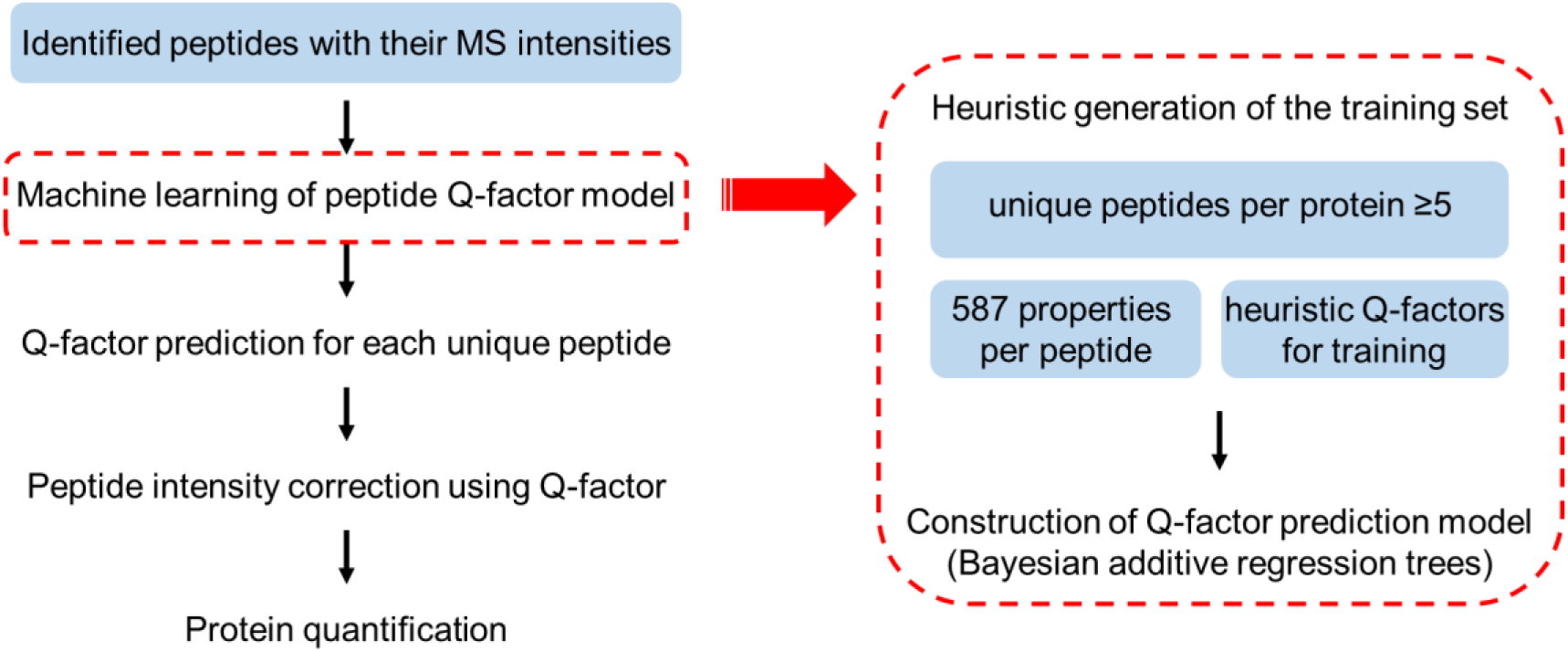
Workflow of LFAQ algorithm. First, based on peptide identifications and MS intensities, the training set of the Q-factor prediction model is automatically constructed using the identified peptides in a complex sample. Proteins with fewer than five identified unique peptides are removed. All the peptides of the remaining proteins are represented by 587 numerical properties. The peptide Q-factors used in the training set are heuristically generated. Then, the BART algorithm is performed to obtain a prediction model of the peptide Q-factor. Finally, the Q-factors of all the identified peptides in the sample are predicted and used to correct their peptide intensities, from which the protein intensities are calculated. The data items are displayed in light blue boxes, and the computational operations are displayed in text boxes.

where N is the number of identified peptides of the protein, and *I*_*i*_ and *Q*_*i*_indicate the raw intensity and the predicted Q-factor of the *i-th* identified peptide of the protein, respectively. It should be noted that the peptides used in LFAQ all refer to unique peptides.

Here, the peptide Q-factor is defined as the extent to which a peptide is quantified by MS (a value between 0 and 1). In other words, it is the ratio of the observed abundance of a peptide expressed by its MS intensity to its actual abundance. For example, a Q-factor of 0.3 indicates that the observed MS intensity of a peptide can reflect only 30% of its actual abundance, and the MS intensity of this peptide should be divided by 0.3 to reflect its actual abundance.

### Peptide Q-factor learning procedure

As shown in **Fig. 1**, the Q-factors of peptides are predicted from a mathematical model built on 587 properties of peptides. The model is trained using a fraction of the identified proteins in the sample rather than spike-in standards. The Q-factor learning procedure can be summarized as follows:

1. Feature calculation: a total of 587 features^14, 17-21^ (**Supplementary Table 1**) describing the peptide sequence and its physiochemical properties are calculated for each identified peptide.
2. Automatic generation of the training set: a heuristic approach is used to generate the training data from the identified proteins in a sample. The training peptides are first collected from the proteins identified by five or more unique peptides, and their 587 calculated features are used as independent variables for regression. Then, the Q-factors of these training peptides, which are estimated heuristically based on a prior model, are used as response variables for regression (see **Methods** for details).
3. Training of peptide Q-factor prediction model: the Bayesian Additive Regression Trees (BART)^22^ approach is applied to the training set to construct the Q-factor prediction model. Based on this model, the Q-factors of all the identified peptides in the sample are predicted and used to correct the raw intensities of the peptides for protein quantification.

### Generation of benchmark datasets

To simulate realistic biological matrices for absolute protein quantification, we designed two samples with different complexities, i.e., yeast and mouse protein extracts, spiked separately with 48 UPS2 standard proteins (Proteomics Dynamic Range Standard, Sigma-Aldrich). The protein mixtures from the yeast and mouse samples were each analyzed by MS in triplicate using both DDA and DIA strategies (**Methods and Supplementary Note 1**). The UPS2 standard proteins are a set of 48 human proteins at six different concentrations (8 proteins per concentration) ranging from 0.05 fmol to 5000 fmol. They are currently the largest set of standard protein mixtures available and are commonly used for evaluating absolute quantification methods in many studies^13, 23, 24^.

**Table 1.** Squared Pearson correlation (R^2^) between the calculated intensities and the actual abundances of the internal standard proteins for the DDA and DIA data of three published papers (ref 1-3). Note that (a) there were two biological replicates (bio1 and bio2) in ref 2, and (b) the iBAQ values in DDA datasets were directly obtained from the reanalysis results of MaxQuant v1.5.2.8, while the iBAQ values in the DIA dataset were calculated based on its definition.

| Dataset<sup>a</sup> | Experimental design | Data acquisition mode | MS instrument | LFAQ | iBAQ<sup>b</sup> | Top3 |
| --- | --- | --- | --- | --- | --- | --- |
| ref 1 | *Mouse+UPS2* | DDA | LTQ-Orbitrap Velos | 0.896 | 0.851 | 0.717 |
| ref 2 (bio1) | *E. coli+UPS2* | DDA | LTQ-Orbitrap | 0.949 | 0.934 | 0.909 |
| ref 2 (bio2) | *E. coli+UPS2* | DDA | LTQ-Orbitrap | 0.956 | 0.927 | 0.915 |
| ref 3 | *E. coli+UPS2* | DIA (SWATH) | TripleTOF 5600 | 0.888 | 0.863 | 0.800 |

### Evaluation of quantification accuracy

To evaluate the quantification accuracy of LFAQ, the three most highly cited absolute quantification algorithms (iBAQ^13^, Top3^12^ and APEX^17^) were used for comparison (**Fig. 2a**, see also **Supplementary Notes 2** and **3**). All four algorithms were evaluated on the same data without additional processes (**Supplementary Table 2**).

**Figure 2.**
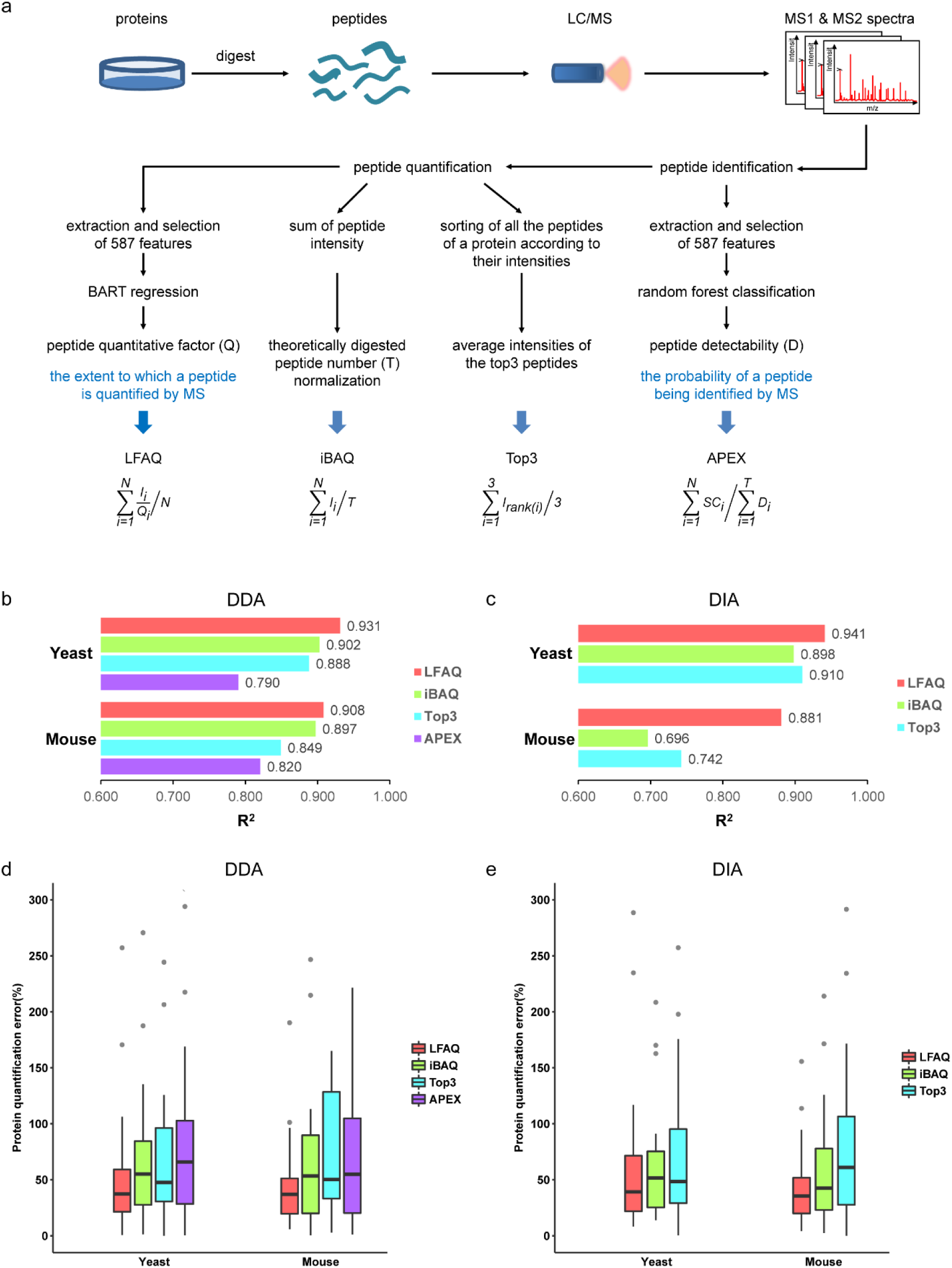
Comparison of different algorithms for label-free absolute protein quantification. (**a**) Illustration of LFAQ and the other compared algorithms. In this study, LFAQ was compared with iBAQ, Top3 and APEX for DDA data and with iBAQ and Top3 for DIA data. In the equations, *N* is the number of the identified peptides of a protein; *T* is the theoretical number of digested peptides of a protein; *Q*_*i*_is the peptide Q-factor of the *i-th* identified peptide of the protein; *I*_*i*_is the raw intensity of the *i-th* identified peptide; and *SC*_*i*_is the spectral count of the *i-th* identified peptide. (**b-c**) The squared Pearson correlation (R^2^) between the calculated intensities and the actual abundances of the spike-in standard proteins in the DDA and DIA datasets obtained using LFAQ and other methods. (**d-e**) Quantification errors (i.e., the quantification errors of the absolute protein abundances) of the UPS2 proteins in the DDA and DIA datasets. In all boxplots, the center black line is the median of the protein quantification errors; the box limits are the upper and lower quartiles; the whiskers delimit the most extreme data points within 1.5 interquartile range below the first quartile and above the third quartile, respectively.

First, we evaluated the correlation between the calculated intensities and the actual abundances of the standard proteins spiked into each sample. As shown in **Fig. 2b-c**, LFAQ showed a higher correlation than iBAQ, Top3 and APEX with at most a 26% increase in the squared Pearson correlation (R^2^). Second, the protein intensities calculated by each algorithm were transformed into absolute abundances using a linear function fitted to the spike-in standards (**Methods**). The quantification errors between the calculated and actual absolute abundances of the UPS2 standards in each dataset were obtained to evaluate the quantification accuracy of these algorithms (**Methods**). As shown in **Fig. 2d-e**, LFAQ displayed smaller quantification error distributions than the other methods with a reduction in error by 24%-29% on the DDA data and 21%-47% on the DIA data, compared with the results of iBAQ and Top3. More importantly, for the low-abundance proteins, i.e., the UPS2 proteins with the lowest identified concentration (5 fmol/L) in this study, LFAQ reduced the quantification error by an average of 46%. These results indicate that LFAQ is more accurate than the other algorithms on all the DDA and DIA datasets.

### Coefficient of variation (CV) evaluation of all the peptide intensities within a protein

The CV distributions of the intensities of all the identified peptides within each protein are shown in **Fig. 3**. The peptide intensity CVs of the proteins with five or more identified unique peptides (i.e., the proteins used in Q-factor training) were markedly reduced after Q-factor correction. Importantly, the same trend was observed for the proteins with fewer than five identified unique peptides, which were not used for Q-factor training. This result implies that the peptide Q-factors were effectively predicted and were not overfitted to the training data. Overall, the median CV of peptide intensities within each protein after Q-factor correction was reduced to less than 5% for all the DDA and DIA datasets, with an average decrease of 39% (DDA) and 77% (DIA) compared to that of the uncorrected MS intensities.

**Figure 3.**
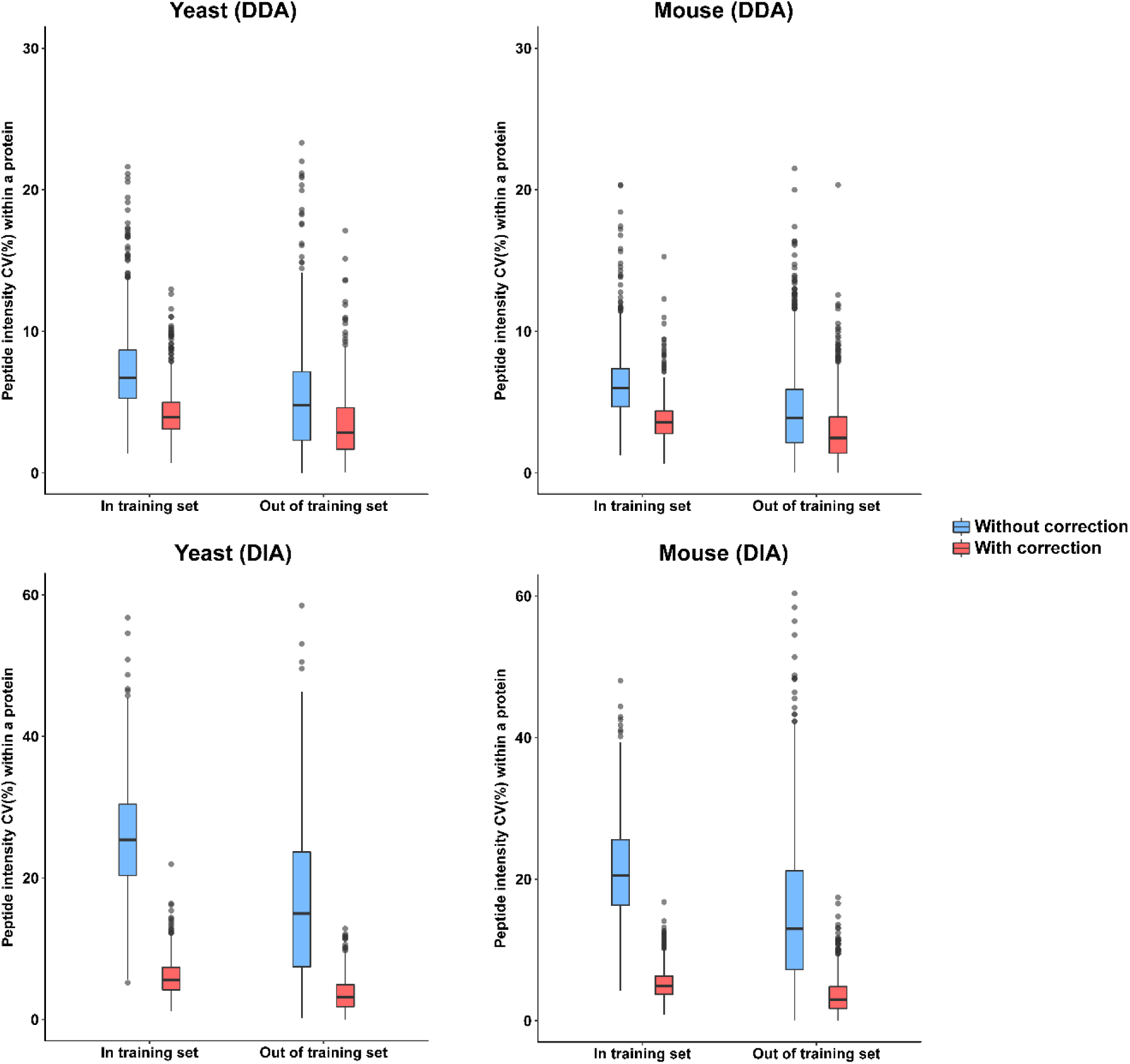
Variance of peptide intensities with and without Q-factor correction in each protein. All the identified yeast and mouse proteins in the DDA (**a-b**) and DIA (**c-d**) datasets are classified into two groups according to whether each protein is used in the Q-factor training set, i.e., whether the identified unique peptide number of a protein is at least five. Proteins with five or more identified unique peptides are used for Q-factor training in LFAQ and are thus classified into the “In training set” group. The “Out of training set” group contains the proteins with fewer than five identified unique peptides, which are not used for Q-factor training. The red and blue boxes represent the CVs of peptide intensities with and without Q-factor correction, respectively. The uncorrected intensity of a peptide was normalized by the molecular weight (Da) of the peptide before CV calculation. Proteins with only one identified unique peptide were not considered here because their peptide intensity CVs cannot be calculated. In all boxplots, the center black line is the median of the protein quantification errors; the box limits are the upper and lower quartiles; the whiskers delimit the most extreme data points within 1.5 interquartile range below the first quartile and above the third quartile, respectively.

### Evaluation of quantification precision

In addition to accuracy, precision is another important aspect in evaluating a quantification algorithm. Quantification precision measures the variance of the quantification results in multiple replicates. Here, the CV distributions of the intensities of the yeast and mouse proteins commonly-identified in triplicate are shown in **Supplementary Fig. 1**. In all the DDA and DIA datasets, LFAQ resulted in a smaller CV distribution than iBAQ and Top3. Moreover, the datasets from human embryonic kidney (HEK) 293 cells analyzed by two MS instruments (Thermo Orbitrap Fusion and Q Exactive HF) in quadruplicate were also used for precision evaluation. The protein intensity CV distributions of the four replicates in both MS instruments further confirmed the precision of LFAQ (**Supplementary Fig. 2 and Supplementary Notes 4** and **5**).

### Validation of LFAQ using public data

In addition to the yeast and mouse datasets we produced in this study, the LFAQ algorithm was also validated on public DDA and DIA data^13, 23, 25^. We reanalyzed the raw data from the papers with the originally described parameters. As shown in **Table 1**, LFAQ displayed higher correlations between the calculated intensities and the actual abundances of the spike-in standard proteins than iBAQ and Top3 in all four public datasets, indicating that LFAQ is an accurate absolute protein quantification algorithm independent of experimental design and sampling conditions.

Furthermore, in this study, the known stoichiometric information on the human 20S proteasome complex^26, 27^ was used to evaluate the accuracy and precision of LFAQ and the other absolute quantification algorithms (**Fig. 5a** and **Methods**). The normalized intensity of each constitutive subunit (α1 to α7, β3, β4, β6, β7) in the 20S proteasome was calculated following the method described in previous papers^26, 27^ (see **Methods** for details). As shown in **Fig. 5b**, LFAQ produced a closer distribution of the normalized subunit intensities to the expected value than iBAQ and Top3 using the data from the paper by Wilhelm M *et al.*^28^ (i.e., the salivary gland and lymph node samples). In addition, LFAQ produced a smaller CV for the 20S constitutive subunits and the three catalytic subunits (β1, β2, β5) with their corresponding immunoproteasome counterparts (β1i, β2i, β5i) (**Fig. 5c**). These results further confirmed the high accuracy and precision of LFAQ.

## Discussion

In summary, LFAQ is the first algorithm for label-free absolute protein quantification, which eliminates the intrinsic biases in MS intensities for all identified peptides using peptide quantitative factor (Q-factor). Its superiority to existing methods in terms of quantification accuracy and precision has been demonstrated on both our own and public data. On average, LFAQ reduced the quantification error by 21%-47% compared to the commonly used methods iBAQ and Top3 on DDA and DIA data. Most importantly, for low-abundance proteins, which are currently the most difficult category to quantify accurately in proteomics, LFAQ reduced the quantification error by an average of 46%.

LFAQ is applicable to any large-scale proteomics experiment due to the advantages of the Q-factor learning algorithm in LFAQ. First, the peptide Q-factor model is trained on the identified proteins in realistic experiments and is independent of spike-in standards. In this study, the known abundances of spike-in standards were used only to evaluate the quantification accuracy of LFAQ and were not involved in the peptide Q-factor learning procedure. Therefore, the superior accuracy of LFAQ was not a result of overfitting to the spike-in standards. Second, the Q-factor model is sample-adaptive, as it is trained from the sample being quantified.

The peptide Q-factor differs from the well-studied peptide detectability in both definition and usage. The peptide Q-factor is directly proposed for peptide quantification and used to correct the biased MS intensity of every peptide rather than only a group of selected peptides, such as the proteotypic or quantotypic peptides. The Q-factor is also different from peptide detectability in terms of algorithm. Unlike peptide detectability learning, which can be handled as a pattern classification problem, peptide Q-factor learning is a multivariate regression problem. Thus, the methods of generating training set also differ. In the Q-factor learning procedure, the training data are automatically generated using a heuristic method without manual curation (**Methods**), based on the fact that the abundances of the identified peptides within a protein should be equal if all the peptides have been fully digested. This approach is essentially different from peptide detectability learning where the training peptides are straightforward to obtain (i.e., a peptide is either identified or unidentified). We compared the effects of the peptide Q-factor and peptide detectability on protein quantification accuracy using the two DDA datasets in this study. As shown in **Supplementary Note 6 and Supplementary Fig. 3**, the peptide Q-factor led to higher quantification accuracy than the peptide detectability.

Finally, we provide an efficient open-source tool with user-friendly graphical interfaces to perform label-free absolute protein quantification on both DDA and DIA data. In theory, LFAQ can use peptide identifications and MS intensities from any software, as it requires only a few essential inputs (**Supplementary Table 3**). Currently, LFAQ supports MaxQuant^29^ (v1.5.2.8 and above) and the public mzQuantML format^30^ (proposed by the Human Proteome Organization Proteomics Standards Initiative) for DDA data and supports SWATH 2.0 (implemented in PeakView^TM^ Software, SCIEX) for DIA data. More tools and formats will be supported in the future.

## Methods

### Preparation and liquid chromatography tandem mass spectrometry (LC-MS/MS) analysis of the yeast and mouse samples

First, samples from Saccharomyces cerevisiae strain BY4743 and Mus musculus RAW264.7 cells were prepared using the methods described in **Supplementary Note 1**. The *Mus musculus* RAW264.7 cells obtained from the American Type Culture Collection were verified by flow cytometric analysis and tested to exclude *Mycoplasma* contamination before experimental use.

Second, UPS2 proteins (1 µg) were digested and added to the yeast and mouse cell digestion mixtures (0.7 µg each). The two samples were then subjected to LC-MS/MS analysis in triplicate using the DDA and DIA strategies separately. The digested peptide mixtures were directly analyzed using a Q Exactive Orbitrap mass spectrometer (Thermo Fisher Scientific) in DDA mode and a TripleTOF 6600 mass spectrometer (SCIEX) in DIA mode. Detailed information can be found in **Supplementary Note 1**.

### DDA and DIA data analysis of the yeast and mouse MS data

For the DDA data produced using the Q Exactive, all raw MS data files were loaded into the MaxQuant workflow (v1.5.2.8). All raw data were searched by Andromeda^31^ against the Swiss-Prot yeast database (release 2013_11) or the mouse database (release 2014_11) with 48 UPS2 standard protein sequences. The decoy sequences were derived by reversing the target sequences at the peptide level^32^, and the search parameters were as follows: (i) an initial precursor ion peak mass tolerance of 20 parts per million (ppm), (ii) a product ion mass tolerance of 0.5 Da, (iii) trypsin enzyme with restrictions on lysine and arginine following proline, (iv) up to two allowable missed cleavages, (v) cysteine carbamidomethylation (+57.0215 Da) as a fixed modification, and (vi) methionine oxidation (+15.9949 Da) as a variable modification. Peptides longer than six amino acids were retained, and both the peptide and protein false discovery rates (FDRs) for the identification results were set to 1%. The iBAQ values were directly calculated in MaxQuant using the default parameters, and the peptide intensities calculated by MaxQuant were used as the input for the LFAQ algorithm.

For the DIA data produced using TripleTOF 6600, the spectral library was first constructed by searching the corresponding DDA WIFF files in the ProteinPilot^TM^ software 5.0 with the same search parameters described above. The resulting group file was then directly loaded into PeakView 2.1 with retention time calibration. The peptide MS intensities were calculated in PeakView 2.1 and the SWATH 2.0 plug-in MS/MS (ALL) with the SWATH Acquisition MicroApp 2.0 software. The main parameters were as follows: (1) the number of peptides per protein was set to 1000, (2) the FDR threshold was set to 1%, and (3) the extracted ion chromatogram (XIC) extraction window was set to 15 min with a 50 ppm XIC width. The results were exported to Microsoft Excel files using the option Quantitation → SWATH processing → Export → All. The “Area-peptides” data sheet in each Excel file was then saved as a separate CSV file and loaded into the LFAQ software for further processing. The iBAQ and Top3 values for DIA data were calculated in the LFAQ software using the equations described in **Fig. 2a**.

### Peptide Q-factor model training

A machine learning algorithm was developed to train the Q-factor model using the identified peptides in the sample, rather than spike-in standards or manually curated data (**Fig. 1**). All the identified unique peptides from the proteins with five or more identified unique peptides are collected as training peptides. For each training peptide, a total of features describing its sequence and physiochemical properties are generated. These features serve as the independent variables (*x*) of the training data.

A key point of the Q-factor learning algorithm is the automatic generation of the training *Q*-factors of peptides, i.e., the response variables (*y*) in the training data. For this purpose, a heuristic method was implemented as described below. Note that the *Q*-factors generated at this stage are used for training only and are not used to correct the peptide intensities for the subsequent protein quantification.

First, peptides from the same protein are assumed to have equal abundances in the sample. Thus, in the training set, the Q-factors of peptides belonging to the same protein should be directly proportional to their observed MS intensities. This assumption is used only for the *Q*-factor training and not for prediction. With this assumption, we need only to determine the *Q*-factor of one peptide within a protein, and the others can be derived from this one. Here, we choose to estimate the *Q*-factor of the most abundant peptide within a protein at first, and the *Q*-factors of the other peptides from the same protein are then calculated as follows:

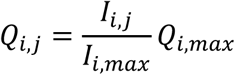

where *I_i,j_* is the MS intensity of peptide *j* of protein *i*; *I*_*i,max*_is the maximum intensity of all the peptides of protein *i* (i.e., the intensity of the top1 peptide of protein *i*); and 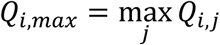 is the corresponding maximum *Q*-factor.

Thus, *Q*_*i,max*_ is the only unknown quantity in the above formula. To estimate *Q*_*i,max*_, we devise here a prior model for the *Q*-factor that is independent of the peptide properties. In this prior model, we regard the *Q*-factor as a random variable and assume that it follows an exponential distribution (see **Supplementary Fig. 4** for empirical evidence), with the expectation θ as the only parameter. Based on this prior model of the Q-factor, we can derive the distribution of the maximum value among a group of Q-factors as follows. Suppose *X*_1_, *X*_2_,…, X_*n*_are *n* independent variables following the exponential distribution of the *Q*-factor. Let *X*_(1)_, *X*_(2)_, …, *X*_(*n*)_be their ordered statistics in ascending order, in which 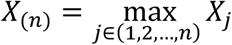. Then, we use the mathematical expectation (mean) of *X*_(*n*)_to estimate *X*_(*n*)_. Below, we briefly describe the procedure to derive the expression of this expectation. Let *W*_1_= *nX*_(1)_and

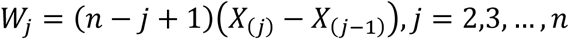

**Figure 4.**
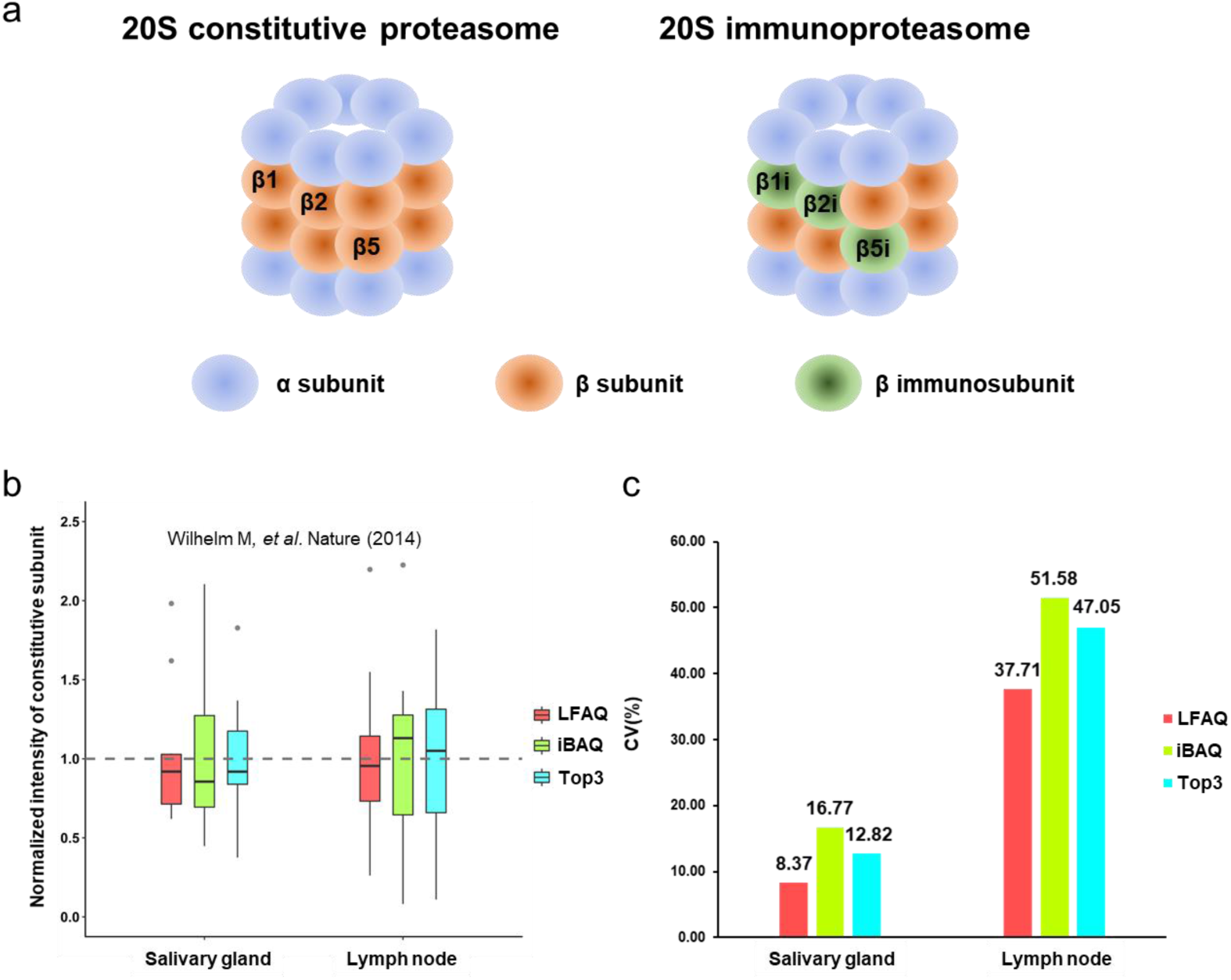
Absolute quantification evaluation of human 20S proteasome. **(a)**Illustration of two main kinds of human 20S proteasome, i.e., the constitutive proteasome and the immunoproteasome. Both proteasomes contain two kinds of subunits (α1-7 and β1-7). There are 11 constitutive subunits (α1 to α7, β3, β4, β6, β7) and three catalytic subunits (β1, β2, β5 and their corresponding immunoproteasome subunits β1i, β2i, β5i) in the 20S proteasome. **(b)** The distributions of the normalized intensities of all the constitutive subunits in human salivary gland and lymph node samples. The theoretical value of the normalized subunit intensities is 1. In all boxplots, the center black line is the median of the protein quantification errors; the box limits are the upper and lower quartiles; the whiskers delimit the most extreme data points within 1.5 interquartile range below the first quartile and above the third quartile, respectively. (**c**) The intensity CV (%) of the 20S constitutive subunits and the three catalytic subunits (β1, β2, β5) with their corresponding immunoproteasome counterparts (β1i, β2i, β5i) in human salivary gland and lymph node samples. Note that the MS data are from the paper of Wilhelm *et al*.^28^.

Then,

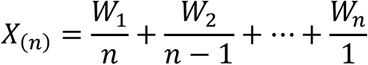

*W*_1_, *W*_2_,…, *W_n_* can be proved to be independent variables following the same exponential distribution as *X_j_* with expectation θ. Therefore, the expectation of *X*_(*n*)_is

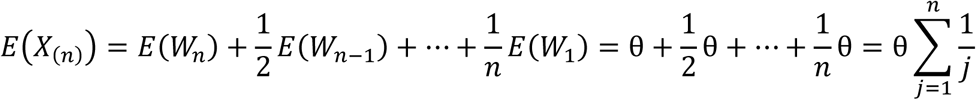

Since the peptides within a protein can be considered independent, the expectation of *Q*_*i,max*_is calculated as follows according to the above result.

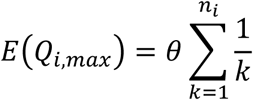

where *n_i_* is the expected number of peptides that can be identified for protein *i*.

Finally, we use the above expectation to estimate *Q*_*i,max*_ and normalize it to the [0,1] interval for all proteins:

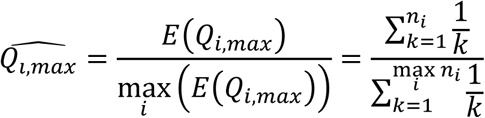

Note that the parameter *θ* is eliminated after normalization and is thus not required to be known.

In practice, *n_i_* is estimated from the length of protein *i*. Note that it cannot be the number of identified peptides in protein *i*, which is affected by the protein concentration in the sample.

In this study, there were 7072 and 9972 training peptides for the yeast and mouse DDA datasets, respectively, and 7182 and 9931 training peptides for the yeast and mouse DIA datasets, respectively (**Supplementary Table 4**).

### Absolute abundance calculation

The protein intensities calculated using LFAQ and the other algorithms can be converted to absolute abundances if internal standards were spiked into the sample. In this study, a linear regression function was used to fit the log-transformed intensities to the absolute abundances of the standard proteins (such as the UPS2 proteins in this study). The absolute abundances of all the identified proteins in the sample were determined from this linear function. We then assessed the confidence of the fitted model by calculating the means and standard deviations of the slope and intersection in the calibration function by bootstrapping (*n*=10,000) under 68% confidence intervals. Detailed information regarding bootstrapping by each quantification method for the DDA and DIA datasets can be found in **Supplementary Tables 5** and **6**. If the number of cells in the sample was known, the protein copy number per cell could be obtained by multiplying the absolute abundance of the protein by the Avogadro constant and dividing the product by the total cell number.

### Relative error of the absolute abundance calculation

In our study, the relative error between the calculated and actual absolute abundances of a protein is defined as:

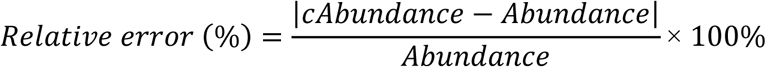

where *cAbundance* and *Abundance* represent the calculated and actual absolute abundances of the protein, respectively.

### Stoichiometric analysis of human 20S proteasome subunits

The human 20S proteasome complex consists of the constitutive subunits (α1 to α7, β3, β4, β6, β7) and the catalytic subunits (β1, β2, β5 and their corresponding immunoproteasome subunits β1i, β2i, β5i). The normalized intensity of each constitutive subunit is defined as follows:

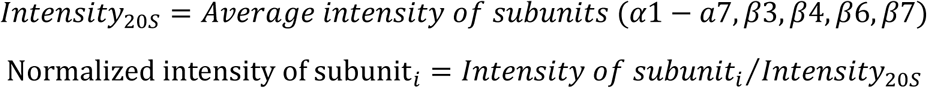

The normalized intensity distribution of all the constitutive subunits was reported to have an expected value of 1^26, 27^ and thus can be used to evaluate the accuracy of the absolute quantification algorithms in this study.

Meanwhile, theoretically, the summed amounts of each catalytic subunit (β1, β2, β5) and its corresponding immunoproteasome subunit (β1i, β2i, β5i) are equal to the average intensity of all the constitutive subunits (α1 to α7, β3, β4, β6, β7)^26^, which can be expressed as follows. Thus, the intensity CV of the constitutive subunit and the three catalytic subunits can be used to evaluate the precision of the absolute quantification algorithms.

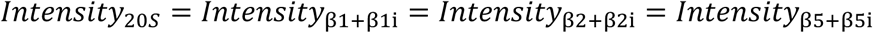

The raw data from the salivary gland and lymph node samples in the paper of Wilhelm *et al.^28^* were downloaded, reanalyzed by MaxQuant (v1.5.2.8). The data were searched against the UniProt human database (release 2014_05). All the other parameters remained unchanged compared to the previous paper. The normalized intensities of all the 20S subunits were calculated *vide supra* using the LFAQ, iBAQ and Top3 algorithms. The Top3 value was first normalized by the protein molecular weight (kDa) for further analysis.

## Code availability

All the source codes and the compiled software are available upon request.

## Data availability

All the raw MS data produced in this study are public available in the ProteomeXchange Consortium via the iProX partner repository (http://www.iprox.org) with the identifiers PXD009719 and PXD009720.

## Acknowledgments

We acknowledge Prof. Li Tang (Beijing Proteome Research Center, Beijing, China) for providing yeast samples, Prof. Matthias Selbach (Max Delbrück Center for Molecular Medicine, Berlin, Germany) for providing MS data, Prof. Henning Hermjakob (European Bioinformatics Institute, Hinxton, UK) and Prof. Haixu Tang (Indiana University, Bloomington, IN, USA) for their helpful discussions. This work was supported by the National Basic Research Program of China (2017YFA0505002, 2017YFC0906602, 2016YFA0501300 and 2014CBA02001), the Strategic Priority Research Program of CAS (XDB13040600), the National High Technology Research and Development Program of China (2015AA020108), the International Science & Technology Cooperation Program of China (2014DFB30010), the National Natural Science Foundation of China (21605159 and 21475150), the Innovation Program (16CXZ027) and the NCMIS CAS.

## Author contributions

X.Q., Y. Zhu. and F.H. designed the study. C.C., Y.F. and Z.G. constructed the algorithm. Z.G. and C.C. programmed the software and analyzed the data. Y. Zhao. and G.W. performed the experiments and obtained the MS data. S.W., W.Y., and M.L. provided experimental and analytical insights. C.C., Y.F., W.Y, X.Q. and Y. Zhu. wrote the initial manuscript. All the authors helped edit the final manuscript.

## References

1. Schubert, O.T., Rost, H.L., Collins, B.C., Rosenberger, G. & Aebersold, R. Quantitative proteomics: challenges and opportunities in basic and applied research. Nat Protoc 12, 1289–1294 (2017).

2. Domon, B. & Aebersold, R. Options and considerations when selecting a quantitative proteomics strategy. Nat Biotechnol 28, 710–721 (2010).

3. Altelaar, A.F., Munoz, J. & Heck, A.J. Next-generation proteomics: towards an integrative view of proteome dynamics. Nat Rev Genet 14, 35–48 (2013).

4. Beynon, R.J., Doherty, M.K., Pratt, J.M. & Gaskell, S.J. Multiplexed absolute quantification in proteomics using artificial QCAT proteins of concatenated signature peptides. Nat Methods 2, 587–589 (2005).

5. Brun, V. et al. Isotope-labeled protein standards: toward absolute quantitative proteomics. Mol Cell Proteomics 6, 2139–2149 (2007).

6. Picotti, P., Bodenmiller, B., Mueller, L.N., Domon, B. & Aebersold, R. Full dynamic range proteome analysis of S. cerevisiae by targeted proteomics. Cell 138, 795–806 (2009).

7. Malmstrom, J. et al. Proteome-wide cellular protein concentrations of the human pathogen Leptospira interrogans. Nature 460, 762–765 (2009).

8. Worboys, J.D., Sinclair, J., Yuan, Y. & Jorgensen, C. Systematic evaluation of quantotypic peptides for targeted analysis of the human kinome. Nat Methods 11, 1041–1044 (2014).

9. Bantscheff, M., Lemeer, S., Savitski, M.M. & Kuster, B. Quantitative mass spectrometry in proteomics: critical review update from 2007 to the present. Anal Bioanal Chem 404, 939–965 (2012).

10. Chang, C. et al. Quantitative and In-Depth Survey of the Isotopic Abundance Distribution Errors in Shotgun Proteomics. Anal Chem 88, 6844–6851 (2016).

11. Vogel, C. & Marcotte, E.M. Absolute abundance for the masses. Nat Biotechnol 27, 825–826 (2009).

12. Silva, J.C., Gorenstein, M.V., Li, G.Z., Vissers, J.P. & Geromanos, S.J. Absolute quantification of proteins by LCMSE: a virtue of parallel MS acquisition. Mol Cell Proteomics 5, 144–156 (2006).

13. Schwanhausser, B. et al. Global quantification of mammalian gene expression control. Nature 473, 337–342 (2011).

14. Tang, H. et al. A computational approach toward label-free protein quantification using predicted peptide detectability. Bioinformatics 22, e481–488 (2006).

15. Jarnuczak, A.F. et al. Analysis of Intrinsic Peptide Detectability via Integrated Label-Free and SRM-Based Absolute Quantitative Proteomics. J Proteome Res 15, 2945–2959 (2016).

16. Li, Y.F., Arnold, R.J., Tang, H. & Radivojac, P. The importance of peptide detectability for protein identification, quantification, and experiment design in MS/MS proteomics. J Proteome Res 9, 6288–6297 (2010).

17. Lu, P., Vogel, C., Wang, R., Yao, X. & Marcotte, E.M. Absolute protein expression profiling estimates the relative contributions of transcriptional and translational regulation. Nat Biotechnol 25, 117–124 (2007).

18. Mallick, P. et al. Computational prediction of proteotypic peptides for quantitative proteomics. Nat Biotechnol 25, 125–131 (2007).

19. Fusaro, V.A., Mani, D.R., Mesirov, J.P. & Carr, S.A. Prediction of high-responding peptides for targeted protein assays by mass spectrometry. Nat Biotechnol 27, 190–198 (2009).

20. Webb-Robertson, B.J. et al. A support vector machine model for the prediction of proteotypic peptides for accurate mass and time proteomics. Bioinformatics 26, 1677–1683 (2010).

21. Eyers, C.E. et al. CONSeQuence: prediction of reference peptides for absolute quantitative proteomics using consensus machine learning approaches. Mol Cell Proteomics 10, M110 003384 (2011).

22. Chipman, H.A., George, E.I. & McCulloch, R.E. BART: Bayesian additive regression trees. 266–298 (2010).

23. Arike, L. et al. Comparison and applications of label-free absolute proteome quantification methods on Escherichia coli. J Proteomics 75, 5437–5448 (2012).

24. Schubert, O.T. et al. Absolute Proteome Composition and Dynamics during Dormancy and Resuscitation of Mycobacterium tuberculosis. Cell Host Microbe 18, 96–108 (2015).

25. Tsou, C.C. et al. DIA-Umpire: comprehensive computational framework for data-independent acquisition proteomics. Nat Methods 12, 258–264 (2015).

26. Fabre, B. et al. Subcellular distribution and dynamics of active proteasome complexes unraveled by a workflow combining in vivo complex cross-linking and quantitative proteomics. Mol Cell Proteomics 12, 687–699 (2013).

27. Fabre, B. et al. Label-free quantitative proteomics reveals the dynamics of proteasome complexes composition and stoichiometry in a wide range of human cell lines. J Proteome Res 13, 3027–3037 (2014).

28. Wilhelm, M. et al. Mass-spectrometry-based draft of the human proteome. Nature 509, 582–587 (2014).

29. Cox, J. & Mann, M. MaxQuant enables high peptide identification rates, individualized p.p.b.-range mass accuracies and proteome-wide protein quantification. Nat Biotechnol 26, 1367–1372 (2008).

30. Walzer, M. et al. The mzQuantML data standard for mass spectrometry-based quantitative studies in proteomics. Mol Cell Proteomics 12, 2332–2340 (2013).

31. Cox, J. et al. Andromeda: a peptide search engine integrated into the MaxQuant environment. J Proteome Res 10, 1794–1805 (2011).

32. Elias, J.E. & Gygi, S.P. Target-decoy search strategy for increased confidence in large-scale protein identifications by mass spectrometry. Nat Methods 4, 207–214 (2007).

## References 1-3

1. Schwanhausser, B. et al. Global quantification of mammalian gene expression control. Nature 473, 337–342 (2011).

2. Arike, L. et al. Comparison and applications of label-free absolute proteome quantification methods on Escherichia coli. J Proteomics 75, 5437–5448 (2012).

3. Tsou, C.C. et al. DIA-Umpire: comprehensive computational framework for data-independent acquisition proteomics. Nat Methods 12, 258–264 (2015).

